# SPArrOW: a flexible, interactive and scalable pipeline for spatial transcriptomics analysis

**DOI:** 10.1101/2024.07.04.601829

**Authors:** Lotte Pollaris, Bavo Vanneste, Benjamin Rombaut, Arne Defauw, Frank Vernaillen, Julien Mortier, Wout Vanhenden, Liesbet Martens, Tinne Thoné, Jean-Francois Hastir, Anna Bujko, Wouter Saelens, Jean-Christophe Marine, Hilde Nelissen, Evelien Van Hamme, Ruth Seurinck, Charlotte L. Scott, Martin Guilliams, Yvan Saeys

**Affiliations:** Data Mining and Modeling for Biomedicine Group, VIB Center for Inflammation Research, Ghent,Belgium; Department of Applied mathematics, Computer Science and Statistics, Ghent University, Ghent,Belgium; Laboratory of Myeloid Cell Biology in tissue homeostasis and regeneration, VIB IRC, Ghent, Belgium; Laboratory of Myeloid Cell Biology in tissue damage and inflammation, VIB IRC, Ghent, Belgium; Department of Biomedical Molecular Biology, Ghent University, Ghent, Belgium; Spatial Catalyst, VIB, Belgium; Laboratory for Molecular Cancer Biology, Center for Cancer Biology, VIB, Leuven, Belgium; Department of Oncology, KU Leuven, Leuven, Belgium; Department of Plant Biotechnology and Bioinformatics, Ghent University, Ghent, Belgium; VIB Center for Plant Systems Biology, Ghent, Belgium

## Abstract

Current spatial transcriptomics technologies are increasingly able to measure large gene panels at subcellular resolution, but a major bottleneck in this rapidly advancing field is the computational analysis and interpretation of the data. To bridge this gap, here we present SPArrOW, a flexible, modular and scalable pipeline for processing spatial transcriptomics data. SPArrOW improves cell segmentation and leads to better overall data quality, resulting in more accurate cell annotations at the single-cell level. Furthermore, it provides the users with numerous visual quality checks that are crucial for the correct interpretation of the data, offering users more control in processing their data. Our workflow is designed to accommodate the various available spatial transcriptomics platforms. Finally, SPArrOW offers interactive visualization and data exploration, enabling sample-specific pipeline optimization by various tuneable parameters and an efficient comparison of different staining and gene allocation strategies.

## Introduction

Recent advances in spatial omics, measuring a multitude of transcriptomic, proteomic, epigenomic^1^ or metabolomic measurements^2^ with an ever-increasing plexity^3^, allows cells to be studied in their native tissue context^4,5^. For spatial transcriptomics, either sequencing or imaging-based readouts can be used, often resulting in a trade-off between the plexity and the resolution of the technology. Some methods such as 10X Visium^6^, MGI STOmics^7^, and Nanostring GeoMX^8^ are untargeted, and thus able to perform whole transcriptome measurements, but measures spots covering multiple cells. Other methods do not allow full transcriptomic measurements but are targeted to a limited panel of genes defined by the user. Next to sequencing-based methods, other technologies offer subcellular resolution and are often based on in situ hybridization (ISH), such as Vizgen MERSCOPE^9^, RNAScope^10^ and Nanostring CosMX^11^. A more elaborate overview of current spatial transcriptomics technologies can be found in literature.^4,12,13^ These latter subcellular resolution methods are particularly promising to study individual cells in their spatial environment, but also pose additional challenges.

Next to challenges in experimental design such as sample preparation and the choice of the marker panel^14^, several computational hurdles remain. Correctly identifying the cellular identity and transcriptomic cell state of each individual cell in the spatial transcriptomics data requires accurate cell segmentation, transcript allocation to cells, and finally cell annotation. ^15,16^ Thus, the technological challenge has moved from the ability to acquire highly multiplexed spatial transcriptomics data to the capacity to efficiently integrate and analyse these datasets. As with all new technologies, there are various options for such analyses but, best practices for processing and quality (QC) control still have to be defined and benchmarked^16^, and we are currently only at the start of this journey^15^.

Existing solutions for cell segmentation highly depend on appropriate staining and face issues such as how to distinguish actual cells from segmentation artefacts. Furthermore, staining images often contain platform specific artefacts (e.g tile borders and illumination gradients, …), requiring preprocessing to optimize the subsequent segmentation. Once a segmentation mask for each cell has been obtained, a feature matrix will be composed by allocating transcripts to a specific cell. In case of nuclei staining, gene allocation can be done in various ways, including restriction to the nuclear mask, expansion with a fixed factor or using the transcriptional composition to determine the cell boundary^17^. There are also various normalization strategies available for the obtained gene count matrix^18^. Finally, once cells have been normalized and clustered, they can be annotated using a variety of methods, ranging from established methods for scRNAseq data to dedicated algorithms for spatial transcriptomics that rely on scRNAseq reference data sets such as Tangram^19^.

Different attempts to create pipelines for spatially resolved transcriptomics that combine all these steps to go from raw input data to annotated cells have been made. Most of the currently available methods do not perform all steps or are not up-to-date with the state-of-the-art building blocks (see supplementary Table 1)^20–27^. Starfish^22^ was one of the earliest computational pipelines developed for imaging-based transcriptomics, but it does not incorporate cell type annotation nor the latest state-of-the-art segmentation algorithms. Furthermore, many of the existing pipelines are not scalable^22,24,27^ and some are not available for all platforms^24,26^. A more recent pipeline, Sopa^20^ focuses on scalability and platform dependence, offering numerous advantages. Nevertheless, it does not include image preprocessing and its annotation capabilities are rather limited. Moreover, as with most other tools (see supplementary table 1), Sopa^20^ also lacks intermediate quality control steps and does not encourage dataset-specific parameter optimization.

To address these shortcomings and meet the demands of current spatial omics technologies, we developed SPArrOW, a SPAtial transcriptOmics Workflow where the basic building blocks are tightly integrated and highly scalable to process spatial omics datasets in a parallelized fashion. SPArrOW stands out by being more comprehensive and interactive, incorporating crucial quality control measures at each step, making it possible to adapt the pipeline to your specific dataset and ensuring more precise and reliable results.

## Results

### SPArrOW offers a scalable, modular and interactive workflow for spatial omics data processing

Our first goal was to ensure the modularity of SPArrOW. Thus, the SPArrOW workflow consists of different building blocks that can be flexibly combined, and that offer different ways of performing every step in the pipeline, depending on the needs of the user. A first building block groups five **image analysis/preprocessing** steps that can be either included or not, depending on the image quality. These include max z-stack projection, illumination correction, inpainting, min-max filtering, and local contrast enhancement (Figure 1a). The second building block performs **cell segmentation**. SPArrOW currently supports segmentation based on watershed, Mesmer^28^, CellPose^29^ and Baysor^19^, and gives the user flexibility in supplying other/custom segmentation models. Segmentation can be done both at the nucleus or cytoplasmic level, offering the possibility to do detailed downstream processing if analyses at the whole cell or only at the nucleus level are required. The next building block performs **transcript allocation**, which can be done either in a naïve fashion, where every transcript is assigned to its location-corresponding segmentation object if there is one or by using different methods of cell expansion, where cells are expanded beyond their segmented body. Depending on the panel size, cell area or library size is used to normalize the counts^18^. Finally, the last building block represents the **cell type annotation** step, where three possibilities are offered: Tangram^19^, labels based on module scores for marker gene sets classically used in scRNAseq analysis^30^ and a built-in SPArrOW annotation optimized for targeted panels based on gene set enrichment scores. For each of these building blocks, SPArrOW uses the SpatialData^31^ object to store all the intermediate information, permitting users to perform the analysis in multiple steps (Figure 1b). Using this platform for data saving and lazy loading makes the analysis more scalable since there is no need to store all intermediate data in memory, nor to load the whole dataset during analysis in memory.

**Figure 1:**
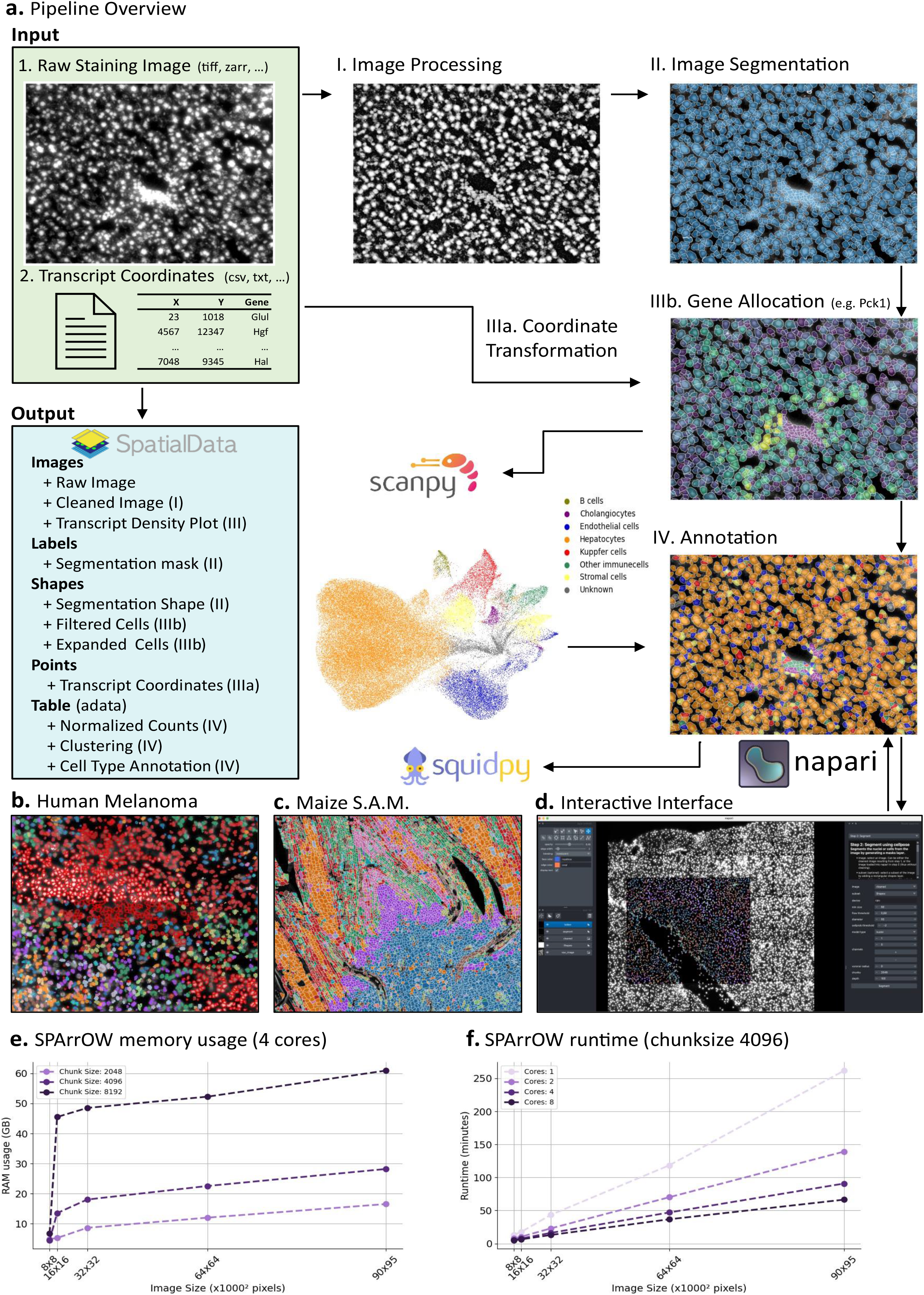
**a:** Schematic overview of the SPArrOW pipeline, demonstrated on a subset of the MERSCOPE mouse liver dataset. Transparent cells with red borders indicate cells that were segmented and subsequently filtered out due to low counts or small size. The output of the various steps is saved on disk in a SpatialData object that enables interoperability with scanpy and squidpy. **b**: Application on human melanoma (Molecular Cartography). **c:** Application on maize shoot apical meristem (S.A.M., Molecular Cartography). **d:** The interactive interface of napari can be used for both pipeline optimization and visualization (demonstrated on mouse liver, Molecular Cartography) **e-f:** The scalability of SPArrOW with the maximum RAM usage summed over all cores **(e)** and the runtime shown in minutes for the whole pipeline, ran on GPU **(f)**.

Our second goal was to build in maximal flexibility for SPArrOW users, setting it apart from existing pipelines. Instead of a single, automatically run pipeline such as SOPA^20^ and Molkart,^24^ SPArrOW permits users to fine-tune specific steps through an optional graphical interface. Users can adjust parameters and explore the impact of including or excluding different steps in the pipeline, via a graphical user interface through Napari (Figure 1c) or by using Jupyter Notebooks. This feature enables users to fine-tune certain steps on a subset of their data and immediately interpret the results. Additionally, SPArrOW provides a command line interface (CLI) that allows the pipeline to be run with fine-tuned parameters on the full dataset, utilizing the Hydra framework.

As spatial datasets can contain images reaching hundreds of gigabytes, we placed a strong emphasis on the scalability of the SPArrOW workflow and implemented this in various ways. Firstly, by utilizing Dask^32^, all image processing steps can be easily parallelized using either CPU or GPU acceleration, where faster runtimes can be achieved with the more available computing power. Secondly, SPArrOW can also be run on lower-end hardware, such as laptops, due to its lazy loading scheme built on Zarr^33^, offering the option to work with larger-than-memory datasets. The flexibility of chunk size allows SPArrOW to adapt to the amount of RAM available. Thus, even users with limited memory can still run the pipeline, although at the expense of running times. Figure 1d shows the scaling behaviour of SPArrOW, both concerning memory usage and computing time. As a concrete example, analysing a Vizgen Merscope dataset of one square centimetre of liver tissue containing 300,000 cells and a panel of 347 genes takes about 1h15 min with eight cores (figure 1f).

Finally, SPArrOW is designed to grant maximal versatility of data type input, both regarding technological platforms as well as different tissue types. SPArrOW allows input data from the following spatial transcriptomics platforms: Vizgen Merscope^9^, 10X Xenium^34^, Resolve Molecular Cartography^35^, MGI STOmics^36^, Nanostring CosMx^11^, SpatialGenomics^37^ and Stellaromics^38^. Furthermore, SPArrOW has been utilised successfully on various tissues already, including mouse and human liver, skin/melanoma, and even maize roots (Figure 1c). SPArrOW has also been designed to be interoperable with current spatial analysis platforms, including Squidpy^21^, and Scanpy^39^ within the scverse^40^.

### SPArrOW improves the segmentation and annotation of liver cells

To showcase the advantages offered by SPArrOW, we now demonstrate how a human-in-the-loop finetuned set of optimized parameters (SPArrOW optimized) compares to a standard processing pipeline using off-the-shelf CellPose^29^ with default parameters and lacking additional image preprocessing (SPArrOW unoptimized, Figure 2a-f). Results were aggregated from four mouse liver RESOLVE datasets. In total, over the four datasets, SPArrOW can retrieve 33% more cells, with on average 31% more transcript counts per cell, and very similar transcript densities. Our optimized pipeline picks up more cells, and these cells are also bigger (higher number of transcripts), while the cell quality represented by the transcript density is comparable for both scenarios (Figure 2a).

**Figure 2:**
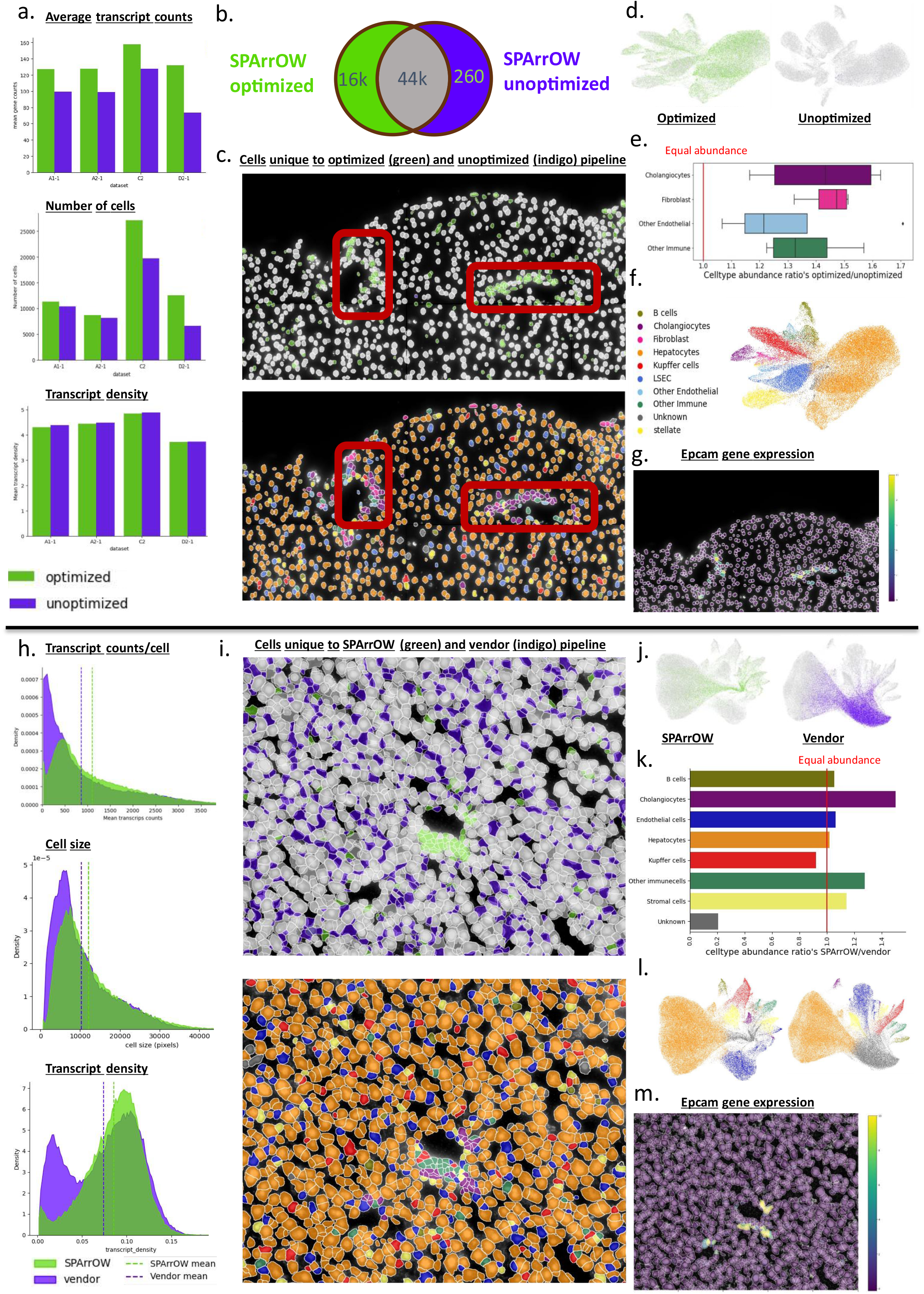
Data quality improvements using SPArrOW. a-g: Comparison of an optimized SPArrOW pipeline versus an unoptimized SPArrOW pipeline (default parameters) across four mouse RESOLVE datasets. **a**: Compares transcript counts, number of cells, and transcript density across the four datasets. **b**. Venn diagram showing matched cells (>50% overlap) between the optimized and unoptimized settings in gray. Green indicates cells only identified with the optimized SPArrOW pipeline, whereas indigo cells are only identified using default parameters. **c**. Zoom-in on slide A1-1. The upper figure colors cells based on their matching status, where green cells are only present when optimizing the pipeline, and blue cells are only present when not optimizing the pipeline (very rare) and the lower figure colors them based on their cell type annotation. Red boxes highlight regions where many cells (green) are only detected with the optimized pipeline. **d** J U ‘ **e**. For a subset of cell types commonly found in vein regions (red boxes in panel c), the ratio of cell type occurrence in the optimized versus unoptimized settings is calculated. The red line indicates equal occurrence. **f**. Combined UMAP of all four optimized slides, colored by cell type. **g**. Gene expression of Epcam, a cholangiocyte marker. h-m Comparison of data analysis using SPArrOW versus vendor (Vizgen MERFISH) on the Vizgen MERFISH mouse liver public dataset. **h**. Distribution plots comparing transcript count, cell size, and transcript density between the two analysis strategies. **i**. Zoom-in view. The upper panel colors cells by matching status, where green cells are only identified when using SPArrOW, and indigo cells are only identified in the vendor analysis. Grey cells are identified in both scenarios. The lower panel shows cells colored by cell type annotation. **j**. UMAP of SPArrOW processed and vendor processed data colored by matching status. **k**. For each cell type, the ratio of occurrence (SPArrOW/vendor) is calculated. The red line indicates equal cell type occurrence. **l**. UMAP of SPArrOW processed and vendor processed data colored by matching status. **m**. Gene expression of Epcam, a cholangiocyte marker.

Interestingly, many of the additional cells that are picked up by the optimized SPArrOW pipeline seem to be region-specific (Figure 2c) as well as cell-type-specific (Figure 2c,d,e). Mainly the portal triad region, enriched in cholangiocytes and fibroblasts suffer from losses when not optimizing the pipeline (figure 2c). When plotting these cells on a UMAP (Figure 2d), the cells only found with the optimized pipeline do not form a separate cluster, and thus likely are not debris or an artefact of the pipeline but rather real cells missed in the unoptimized workflow, demonstrating the power of our pipeline. As an additional quality control check, we assessed the ratio of cell abundance of the optimized versus the unoptimized scenario and found that all cell types are better recovered using the optimized pipeline, but some cell types seem to be impacted more than others (Figure 2e). For example, for cholangiocytes, the optimized scenario results in a 50% increase in the number of cells, which could seriously affect any downstream processing. Indeed, plotting the expression of Epcam a specific cholangiocyte gene further confirmed that the cells in the regions identified by the red boxes in Figure 2c are cholangiocytes (Figure 2g), which might be missed when using default pipelines. Our example thus highlights the importance of careful data processing, and the human interactive finetuning offered by SPArrOW supports such comparative analyses. Furthermore, this again clearly demonstrates that any parameter optimization during data (pre)processing can greatly influence any downstream analysis and conclusions.

While many vendors offer spatial transcriptomics technologies, many of them do not provide state-of-the-art analysis pipelines, and even if they do these solutions lack key functionalities, do not include the latest state-of-the-art methods, are not tailored to your specific dataset and are not broadly applicable. To assess how SPArrOW compares with such pipelines, we compared the analysis performed by the SPArrOW pipeline to the segmentation offered by the Vizgen MERFISH technology (vendor)^9^, supplemented by the SPArrOW steps allocation and annotation (Figure 2h-m).

Comparing the distributions of the transcript counts, cell size and transcript density, demonstrates that SPArrOW picks up fewer cells than the vendor pipeline, but the majority of the additional cells picked up by the vendor pipeline concerns cells with very few transcripts per cell, potentially hinting to more debris being picked up by the vendor pipeline. The cells picked up by SPArrOW also seem to be high-quality cells, represented by a higher transcript density (Figure 2h). The spatial distribution of the cells specifically picked up by one pipeline and not the other is shown in Figure 2i. Again, the cells specifically picked up by SPArrOW seem to be more region and cell type specific, while the vendor-specific cells are everywhere and of low quality (low transcript density). Moreover, they often do not overlap with the polyT staining, most likely indicating that no cell is present in these areas (Figure 2i).

To further characterize the difference in the cells being identified, we performed an automatic cell type annotation (Figure 2i, bottom plot and Figure 2l). From this, we found that the vendor-specific cells (denoted in purple; “ “ in Figure 2l. This can be explained since the cell type classifier cannot reliably predict a cell type due to the very low transcript density of these cells. For the cells specific for the SPArrOW optimized pipeline, we again observe that a large part of them are region and cell-type specific, again enriching the cholangiocytes around the veins, as evidenced by the Epcam expression (Figure 2m). While most cell types are equally represented in both analyses, many more cholangiocytes are retrieved by SPArrOW, and to a lesser degree, this is true for immune cells and stromal cells (Figure 2k). Thus, while SPArrOW picks up 15% less cells, this does not result in less annotated cells. Kupffer cells (KCs) are the only exception to this. Due to their irregular shape and long arms, they are more often missed when segmenting using polyT staining. Taken together, SPArrOW results in an improved annotation: it leads to a lot fewer “ “ cells are returned by the vendor pipeline, and more cells of the other cell types can be annotated than with the results offered by the vendor pipeline.

### SPArrOW facilitates improved quality control and interpretation of spatial omics data

The ability to arrive at high-quality processed data for spatial omics is a crucial, but often overlooked step. A key advantage of SPArrOW is its potential to help the user in more rigorous quality control (QC), and this during the different steps in the analysis pipeline. Compared to existing pipelines, as well as some of the individual building blocks such as image preprocessing or cell segmentation, SPArrOW allows for better QC checks and interpretation, and this in an interactive way at each step of the pipeline. The advantage of intermediate QC checks is that a user can build more confidence in the pipeline results, as it allows them to see where artefacts might be introduced, which is crucial for downstream analysis and interpretation and enables the pipeline to be tuned accordingly, so some artefacts can be removed.

SPArrOW offers several visual quality checks, including assessments of equal cell brightness, background signal levels, artefact spotting, segmentation quality, and cell quality, such as checking if sufficient transcripts are allocated (Figure 3). These checks are linked to user-tunable parameters that enhance image quality or filter out low-quality cells. At each step of the analysis, all intermediate results are saved in the SpatialData object, ensuring no loss of information. Additionally, the built-in annotation tool quantifies the number of cells that cannot be reliably annotated (“unknowns”), as demonstrated in the previous section (Figure 3).

**Figure 3:**
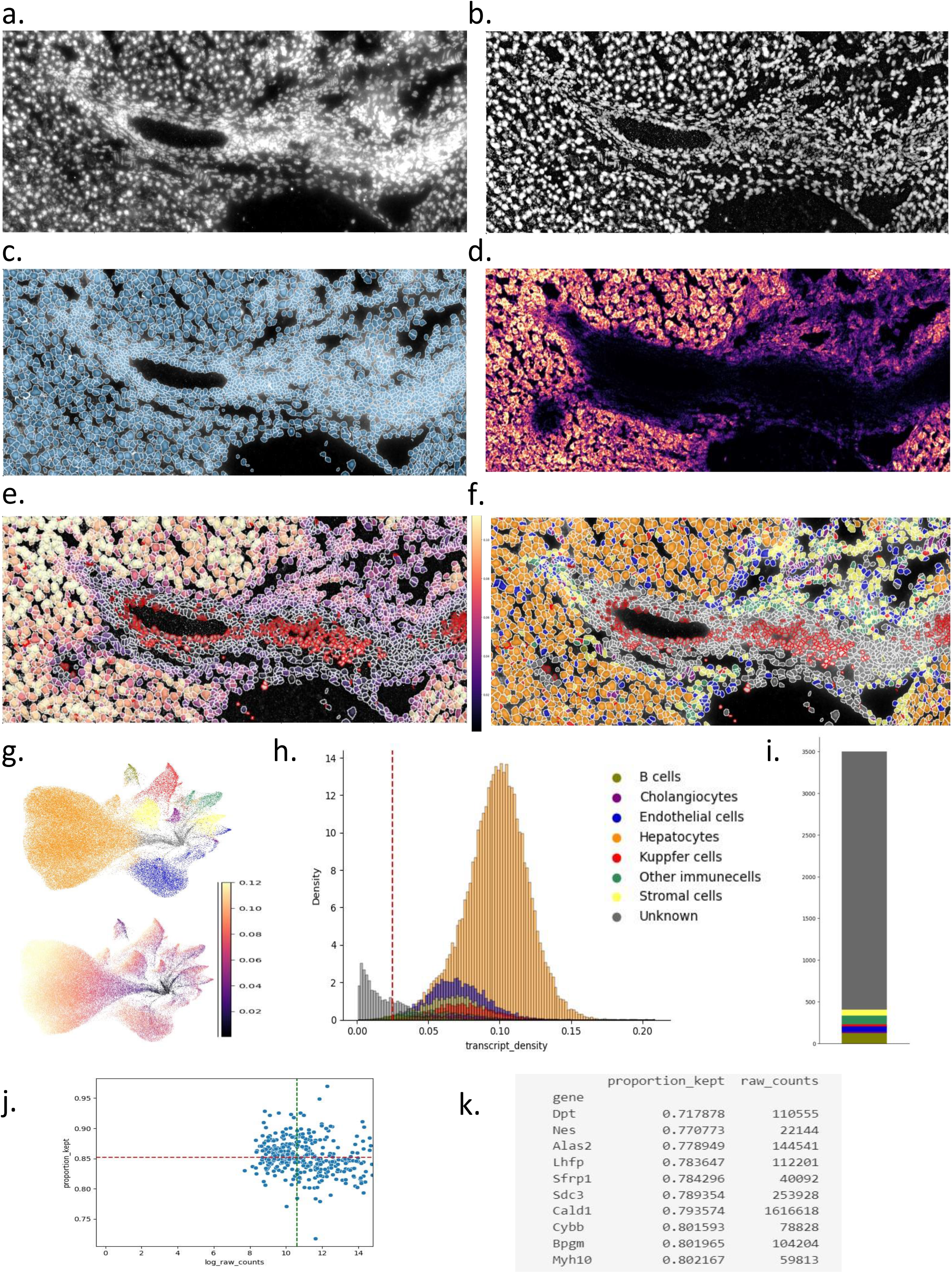
Quality control using SPArrOW. a-e: subset of MERSCOPE mouse liver data. **a**. raw input image **b**. processed image after applying SPArrOW image processing steps **c**. raw images overlayed with segmentation mask. **d**. Transcript density plot created solely based on transcript locations. **e**. Cells colored by transcript density. Cells with a red border indicate those with a transcript count lower than 5, which were filtered out. **f**. Cells colored by celltype annotation. **g**. UMAP’s colored by celltype annotation (Upper), and transcript density (Lower). **h**. Density plot of transcript density split by cell type. The red dotted line indicates the recommended transcript density threshold (0.025 for this dataset). **i**. Celltype composition of cells under the transcript density threshold of 0.025 **j:** Scatter plot showing log transcript counts on the x-axis and the percentage of transcripts present in cells on the y-axis. The red dotted line indicates the average percentage of transcripts retained, and the green dotted line indicates the average expression level of a gene. **k:** Output of SPArrOW: a table indicating the genes that are the least retained in the analysis.

Based solely on the input transcript coordinate file, SPArrOW allows the investigation of transcript density. Specific regions may contain cells but lack transcripts (Figure 3 b-d). Possible reasons for this include oversaturation (e.g., due to insufficient tissue clearing), high necrosis areas, cells lacking marker genes, collagen-rich regions, etc. This can be useful feedback for improving dataset generation in follow-up experiments. The QC plot can also identify areas with high transcript density but without cell staining, potentially indicating staining failures.

Within SPArrOW, cells with fewer than a pre-specified number of transcripts (here, five) and cells that are too large or too small are filtered out. These cells are still visualized with a red border (Figure 3 e-f) to identify region-specific quality issues (around the portal triad) and confirm that the information loss isn’t due to segmentation errors.

The built-in cell type annotation tool in SPArrOW allows cells to be annotated as unknown when insufficient information is present. These cells often have a low transcript density (Figure 3 e-h). Filtering based on transcript density, instead of size and counts, might be more effective. For this dataset, setting the threshold at 0.025 transcripts per pixel filters out cells that are mainly annotated (85%) as unknown (Figure 3 h-i). This threshold, however, is dataset-specific, as a low transcript density may also result from a lack of markers for certain cell types.

When allocating transcripts to cells, not all transcripts will be within segmented cell boundaries due to imperfect stainings, segmentation errors, diffusion, etc., resulting in some transcripts being lost in the allocation step. Quantifying the information loss for each gene helps identify outlier genes with a disproportionately large number of lost counts (Figure 3 j-k). For example, observing many genes here specific to a particular cell type could indicate that staining for that cell type would have worked less well.

## Discussion

Advances in spatial omics technologies have yielded an ever-growing broad variety of different platforms that allow users to measure transcriptomic information at subcellular resolution. However, the diversity of the existing technologies, as well as the large volumes of data that are produced by these platforms necessitate the development of scalable data processing pipelines that are sufficiently flexible to accommodate data from different platforms, tissues and organisms, yet offer a common interface to their data processing.

Here, we introduce SPArrOW, a modular, scalable and interactive workflow that flexibly accommodates spatial transcriptomics data from a variety of platforms. SPArrOW offers a unified pipeline that can be used for all subcellular resolution datasets across platforms, tissues and species, reducing the learning curve for analysing new methods and minimizing bias when comparing quality between techniques. SPArrOW “ “ platform for spatial transcriptomics data analysis would not be compatible with the ongoing evolution in the field, and thus rather providing a modular workflow with flexible finetuning of the different building blocks.

A unique advantage of SPArrOW, compared with existing tools, is its modularity and interactivity, allowing users to perform intermediate parameter tuning and quality control, leading to better data quality and improved downstream processing and interpretation. In this way, SPArrOW embraces the “human in-the-loop”concept, the leading to more trueworthly analysis pipelines for spatial omics data.

Once the user is satisfied with the parameter optimization, SPArrOW allows large-scale processing of arbitrary-sized datasets using parallelization and lazy loading techniques that support”larger than memory”datasets.

Here we have showcased the advantages of SPArrOW in several use cases for liver tissues. By comparing SPArrOW to both classical, unoptimized as well as vendor pipelines, we show that SPArrOW improves segmentation and interpretation of spatial omics data. Parameter finetuning in SPArrOW shows that certain regions- as well as cell-type specific cells, are better recovered using an optimized SPArrOW pipeline and that this leads to improved cell-type annotation as well as biological interpretation of spatial patterns in the liver. Certain cell types in the liver, such as cholangiocytes and fibroblasts are much better recovered using the SPArrOW pipeline compared to existing approaches. Thus, from our study, it is clear that improved quality control and preprocessing of spatial data can have major implications for downstream data analysis and interpretation.

A major addition that SPArrOW brings to the current toolset of spatial omics data processing pipelines is its built-in support for quality control. As highlighted already in many recent works, good QC checks are vital for any (spatial) single-cell pipeline^4,15^, and SPArrOW contributes to this important research direction by offering the user many visual QC checks. These include checks for the quality of segmentation, transcript allocation, and transcript density, but also downstream steps such as cell type annotation can contribute important information. As an example, we demonstrated on liver tissue samples that better quality control largely impacts downstream cell annotation. In addition, many of the new QC plots also shed light on the quality of the original raw input data, e.g. potentially showing staining problems or artefacts.

In summary, we presented SPArrOW, a new modular and scalable workflow that improves spatial transcriptomics data analysis and interpretation and implements a human-in-the-loop strategy to improve data quality control and parameter optimization with support through a graphical user interface.

## Online Methods

### Datasets and technologies

SPArrOW has been run on a plethora of datasets from various platforms (Vizgen Merscope^9^, 10X Xenium^34^, Resolve Molecular Cartography^35^, MGI STOmics^36^, Nanostring CosMx^11^, SpatialGenomics^37^ and Stellaromics^38^). For the analysis shown in this paper, 6 different datasets were used.

First, we used a public mouse liver MERSCOPE (Vizgen) dataset^41^, animal 1 replicate 1. The dataset contains 347 genes and five different cell stainings, including DAPI, polyT, and three other unspecified membrane stainings. Analyses were limited to a subset of the data (45056 by 45056 pixels), except for the scalability analysis, which was performed on the entire section (90,000 by 95,000 pixels) with a pixel size of 108 nanometers. In-house comparisons revealed optimal results for polyT in combination with the DAPI staining compared to the other stainings, which was subsequently used in the rest of the analysis.

Secondly, four slides of mouse liver from RESOLVE molecular Cartography were used (A1-1,A1-2, C2,D2-1) each containing 100 genes and a DAPI staining^42^. The fifth available slide, containing a small part of the capsule, was discarded as the tissue architecture was less comparable to the other sections.

Thirdly, a section of maize shoot apical meristem was used in Figure 1. This dataset was generated using RESOLVE Molecular Cartography and is a pilot study with a limited panel of 20 genes and a cell wall staining (Calcofluor white). The dataset size is 8192*12288 pixels (138 nm per pixel).

Fourthly, a human melanoma slide of 38592 × 34304 pixels was used (slide 5 region A1-1)^43^. The dataset, generated with RESOLVE Molecular Cartography, contained 100 genes and a DAPI staining.

### Pipeline implementation

SPArrOW is implemented in Python with the optional installation of the Julia-based package Baysor^17^. Python was selected due to its extensive libraries for image processing and bioinformatics, scalability, and a growing user community. The pipeline is built on the infrastructure of SpatialData^31^, which is itself based on OME-Zarr^33^, Dask^32^, xarray, geopandas, and Anndata^44^. This framework was chosen for several reasons: it supports lazy loading, which is essential when processing large spatial datasets that cannot be loaded into memory; it integrates seamlessly with the scverse ecosystem^40^; it handles big data efficiently; and it provides storage options for all data types utilized and generated during the SPArrOW workflow.

### Pipeline Scalability

The pipeline’s scalability is enhanced by utilizing Dask^32^ and SpatialData^31^. Dask facilitates parallel execution, larger than memory processing and distributed computation by breaking large datasets into smaller chunks and processing them in parallel, thereby enabling efficient memory management and handling of large-scale data. The interoperability between the SpatialData Zarr format and Dask allows for efficient reading and writing of chunked data to a Zarr store^33^.

The SpatialData object contains the following elements: Images (raster images), Labels (raster segmentation masks), Points (transcripts), Shapes (segmentation boundaries) and Tables (cell counts matrix). Due to the potentially large size of the data, scalable processing on Images, Labels and Points is essential for efficient handling.

Scalable preprocessing of images is implemented in SPArrOW by providing an apply function, that can take almost any image processing function as input, with SPArrOW handling the scaling utilizing Dask. Similar processing on Labels is provided via a dedicated apply function, where care is taken to prevent collisions between labels in different chunks. SPArrOW also implements a segmentation function that can take any callable segmentation model that converts images to masks, with all processing scaled accordingly. For the latter, a minimal overlap between the chunks is required (∼cell size), depending on the algorithm chosen by the user for merging segmentation masks resulting from different chunks. A similar segmentation function is provided that can take any callable function that converts transcripts (Points) to masks. For the allocation of transcripts to cells (Labels/Shapes) and calculation of transcript density, SPArrOW also provides scalable utility functions.

Both Shapes and Tables are handled efficiently in SPArrOW by utilizing the AnnData^44^, Scanpy^39^, GeoPandas and Shapely libraries.

### Accessibility

SPArrOW can be accessed in three different ways, as an interactive tool in Napari ^13^, as a command line interface using Hydra and via the API with tutorials and explanations available in Jupyter Notebooks ^13^.

The Napari^45^ plugin was created using the Napari cookie-cutter template. The plugin allows users to interactively tune parameters for various steps of the pipeline on a data subset in real-time, providing a non-coding solution for biologists to analyze their data, view results, and better understand any limitations. Napari was chosen because it is an open-source, scalable tool with a user-friendly interface that seamlessly integrates with other Python libraries, as it is programmed in Python.

Once parameters are optimized, the CLI using Hydra can be employed to analyze the complete dataset. Hydra was chosen for its flexible configuration management and support, via the Hydra Submitit Launcher plugin, for High Performance Computing.

Another approach involves using the API within a Jupyter Notebook, enabling users to both fine-tune parameters on a subset and subsequently run the full analysis. Example notebooks with parameter explanations are available on the readthedocs page.

### Data Storage

The SpatialData^31^ object stores the following data:

- **Images**: This layer contains all images as separate entities, including both the input images and those generated during the processing pipeline. The data is stored in the OME-Zarr format.
- **Labels**: This layer stores all raster data, specifically the segmentation masks that indicate which pixels belong to which cells.
- **Shapes**: In this layer, all cells are stored as Shapely shapes within a GeoPandas object. This includes objects containing all analyzed cells as well as those with filtered cells. Storing filtered cells makes it possible to visualize them, which helps detect problematic regions. Additionally, shapes of regions of interest can be included in this layer.
- **Points**: This layer contains a Dask data array that holds all transcript locations, enabling precise spatial mapping of transcriptomic data.
- **Table**: This layer consists of an AnnData object that includes gene counts and annotations for each cell, facilitating comprehensive data analysis and integration.

### Pipeline building blocks

#### Image Processing

The image processing steps for SPArrOW are performed separately for each staining channel. These steps can be completed in a single pipeline run with potentially different parameters for each channel.

- **Z-stack projection**: Often, multiple z-stacks of an image are provided. For optimal processing in 2D, a max projection is performed. For each pixel position in the xy-plane, the maximal value over the different z-stacks is kept. SPArrOW advises processing 2D images as 3D segmentation is still in beta.
- **Tiling correction**: To address potential tiling artefacts within the dataset, the BaSiC tool^46^ is used to correct illumination inconsistencies within tiles. It works based on low-rank and sparse decomposition. The tile size is adjustable.
- **Inpainting**: Empty grid lines at the border between tiles can artificially split up cells during segmentation. These lines are removed using OpenCV functionality to ensure the continuity of cell structures. This is done by looking at the neighbouring pixels and ensuring a continuous process, using principles from fluid dynamics^47^.
- **min/max filtering:** Due to autofluorescence or other artefacts, background noise might appear, possibly influencing the segmentation. A min-max filter step can help remove HALOs around cells and eliminate background debris. This is done by looking at the values of neighbouring pixels. The filter size is an optimizable parameter, and filters from Dask image are utilized.
- **CLAHE contrast enhancing**: To achieve uniform illumination across the slide, contrast is enhanced using contrast-limited adaptive histogram equalization (CLAHE). The lightness values are redistributed over separate regions in the image, which is crucial for accurate segmentation.

The results of each step are saved in the SpatialData object by default.

#### Segmentation

As mentioned in the section ‘pipeline scalability’, SPArrOW supports both stain-based and transcripts-based segmentation by scaling any callable that returns masks given images or transcripts. For ease of use, we already provide several options that can be plugged into the segmentation algorithm step to create a mask: Cellpose^29^, watershed segmentation, MESMER^28^ segmentation, and the transcript-based Baysor^17^.

##### Cellpose

Cellpose is the default segmentation tool in SPArrOW due to its robust performance across various datasets and proven track record^48^. Cellpose can segment both nuclei and membrane stainings and supports 3D segmentation. Cellpose is a deep-learning model, with a U-Net backbone^49^. While this method performs well with default settings, tuning certain parameters can significantly improve segmentation quality. The three key parameters to optimize are:

- **Average Cell Size**: Adjusts the expected size of cells. This parameter can be set to None, in which case cellpose estimates the parameter. This estimation, however, is often subpar and requires a lot of computing power.
- **Flow Threshold**: Indicates the roundness of cells. Increasing this parameter allows for fewer round cells to be segmented.
- **Cell Probability Threshold**: Determines the confidence level required for a pixel to be classified as part of a cell.

#### Watershed Segmentation

Watershed segmentation is also available as a baseline method. It is quick and useful for performing preliminary checks but is not recommended for detailed analysis due to its relatively lower accuracy.

#### MESMER Segmentation

MESMER segmentation is another option provided by SPArrOW.

When processing cell wall staining of plant cells, it is often beneficial to color-invert the image to obtain better results.

#### Transcript Allocation

Transcripts are assigned to their respective cells based on the segmentation mask. If the segmentation mask is unavailable or changes were made to the shapes layer, the shapes are converted into new labels layers using the rasterio^50^ library. Using Dask and the segmentation mask, billions of transcripts can easily be assigned to their respective cell, resulting in a cell-by-gene count matrix. In certain cases, cell expansion beyond segmented borders is desirable and can be achieved using a Voronoi diagram as border limits on the shapes or labels layer.

##### Baysor

Baysor offers a segmentation-free approach, deciding on cell borders based on the transcript composition. It can do this without prior segmentation as well as guided by a segmentation mask.

#### Table Processing

The raw count matrix is processed using Scanpy functionality, adapted for targeted spatial transcriptomics data.

Firstly, Cells with low transcript counts (default threshold of five) are filtered out. Genes are excluded if expressed in fewer than 10 cells (default). Filtered cells are saved in a separate object for visualisation purposes. Secondly, cells are normalized. For limited panel sizes, normalization based on cell size is recommended^51^, as transcript counts per cell are heavily dependent on the gene panel choice composition. For large panels and sequencing-based platforms, library size normalization is available.

After size normalization, the cells are log-transformed and scaled. ‘ ^39^ normalize_total function is followed by a log transformation when using library size normalisation. Afterwards, a principal component analysis is performed, followed by UMAP^52^ creation and Leiden clustering^53^.

### Annotation

SPArrOW provides three annotation options, while also allowing the integration of other algorithms via the Anndata object.

- **Tangram**^19^: Commonly used^20^ for annotating targeted spatial transcriptomics data, requiring a matching scRNA-seq atlas. The annotation is performed by mapping the spatial data onto an annotated atlas.
- **Scanpy’s score_genes Function** : The gene panel is split into modules of marker genes per cell type.

The enrichment score for each cell type module is calculated by comparing the average expression for each cell type to other comparable genes in the panel, an algorithm classically used for scoring module enrichment in scRNA-seq data^54^. Every cell is then assigned to the celltype for which it obtained the highest score if this score is higher than zero. However, it may face challenges due to the limited gene panel in targeted spatial transcriptomics data compared to the scRNA-seq data that the tool was designed for.

#### SPArrOW annotation

The built-in annotation algorithm of SPArrOW starts from a gene by celltype matrix ***W*** and the gene expression matrix ***E*** of a sample. Given a gene panel ***G*** and a set of non-filtered cells **C**, a gene *g* is considered a marker for cell type *ct* if the weight w**(g**,**ct)** is positive. The set of marker genes for a cell type is defined as **MG(ct)**:

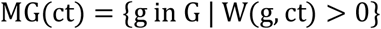

The annotation algorithm calculates a score matrix ***S***_***i***_ in each iteration i. Each row of a score matrix corresponds to a cell in the sample and each column to a cell type in the marker gene list. Before an iiteration for each gene g, a weighted) mean expression vvalue-subscriptalue ***μ***_***i***_(***g***) is calculated over all cells. During an iteration, every cell is scored for every celltype. The weights in the marker gene list are used to give importance to the marker genes. If equal weights are desired, they can be set accordingly. The score of cell c for cell type ct, is the difference between the weighted sum of the expressions of the marker genes in the cell and the weighted sum of the mean expression values of the marker genes in the sample, divided by the number of marker genes.

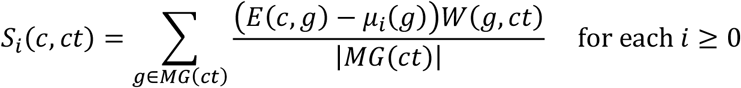

The score matrix ***S***_***i***_ is used to assign a cell type to each cell after iteration. The cell type assigned is the one for which the cell received the highest score. If a cell has no positive score for any cell type, it is annotated as ‘Unknown’. The annotation is temporary since cells can be annotated differently in subsequent iterations. The annotation label of the cells after iteration i is referred to as *label*_*i*_.

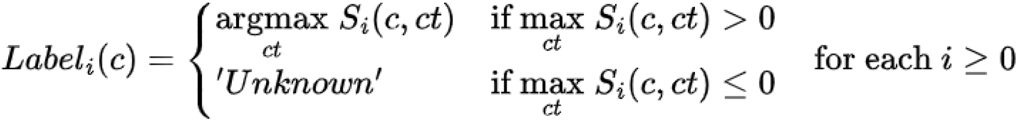

***V***_***i***_(***ct***) is the set of cells annotated as celltype ct in annotation round i. **CT** is the set of all cell types in the marker gene list, and ***CT***_***i***_ is the set of cell types is the set of cell types, excluding ‘Unknown’, present in iteration i.

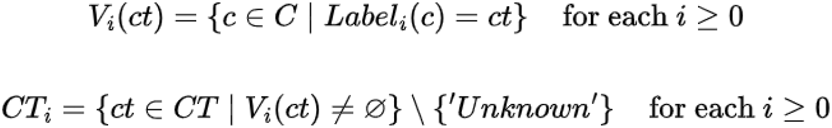

In iteration 0, **μ**_**0**_(***g***) is the average expression of gene g over all the cells. Consequently, the prevalent cell types in the sample influence **μ**_**0**_(***g***) much more than the rare cell types. In iteration i with *i* ≥ 1, this bias is corrected based on the annotation from round i-1. For each cell type ct, except ‘Unknown’, present in annotation i-1, the average expression of the gene g is calculated over the cells annotated as ct. μ_*i*_(*g*) with *i* ≥ 1 is the average of these averaged expressions of g per cell type in annotation round i-1.

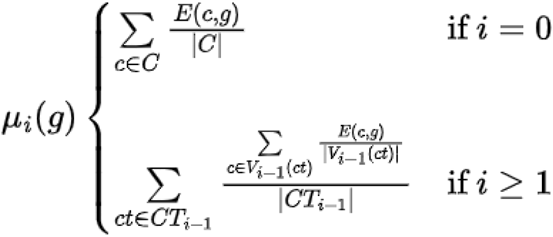

The iterative procedure of the SPArrOW annotation algorithm converges fast. It is stopped after iteration i, if the difference between annotation round i and annotation i-1 is small (e.g.≤ 0.05%) of cells with a different annotation, by default). The annotation algorithm returns the annotation and the final scoring matrix.

The marker gene matrix W can be expert annotated, or derived from the atlas. When deriving W from the atlas, we recommend weighing the marker genes based on a differential expression score, to transfer as much information as possible.

### Cell matching

To compare the same cell segmented in different ways, cells in different samples need to be matched. This is done using geopandas sjoin. Cells are deemed matched if at least 50% of the area of the smallest cell is located in the bigger cell. Multiple cells from one sample can be matched to one cell in another sample (e.g. binucleated cells).

### Transcript density

Transcript density in a cell is calculated by dividing the total transcript counts of a cell by the segmented area.

The transcript density image is created by calculating the number of transcripts measured at every pixel, then adding a Gaussian filter (sigma=7) from the Scipy package^55^.

### Data Availability

Not all steps were included when processing the data obtained by the proprietary analysis performed by Vizgen MERFISH. Complementary to this, no clear overview of the analysis steps was given. We started from the segmented cells the Vizgen analysis provided and performed all the following

## Data Availability

The MERFISH mouse liver data is available at the following website: https://info.Vizgen.com/mouse-liver-data.

The liver RESOLVE Molecular Cartography data is available at the following website: https://www.livercellatlas.org/

## Code Availability

Code availability at the following GitHub page: https://github.com/saeyslab/napari-SPArrOW

