## Supplemental table 1 for "SPArrOW: a flexible, interactive and scalable pipeline for spatial transcriptomics analysis"

### Supplementary Table

| Characteristics | SPArrOW | SOPA | Steinbock | Molkart | STARFISH | PIPEFISH | Squidpy | Giotto suite | Voyager | stereopy |
|---|---|---|---|---|---|---|---|---|---|---|
| Input data | Image + mRNA coord | Image+ mRNA coord | Image+ mRNA coord | Image+ mRNA coord | Fluorescent images | Fluorescent images | Cell*gene matrix | Cell*gene matrix | Cell*gene matrix | GEM/GEF+ opt image |
| Image processing | Yes, tunable | No | Yes | Yes | Yes, tunable | Yes | No | No | No | No |
| Segmentation | Yes, tunable | Yes | Yes | Yes | Yes, watershed | Yes | Yes, external | No | No | Yes |
| Allocation | Yes, tunable | Yes | Yes | Yes | Yes | Yes | No | No | No | Yes |
| Celltype Annotation | Yes, tunable | Tangram | For IMC | No | No | No | Yes, external | No | No | yes |
| Interactive visualization | Yes, Napari | 10x Xenium visualizer | Yes, Napari | No | Yes, Old napari | No | No | Yes, R shiny | No | Yes, in notebook |
| Platform versatility | Yes | Yes | Yes, created for IMC | Molecular Cartography | Yes | Yes | Yes | 10x, MERFISH | yes | stereoSeq |
| GPU accelaration | Yes | Yes | No | No | No | Yes | External | No | No | yes |
| Data Format | SpatialData | SpatialData | Spatial Experiment | Anndata | spaceTx | spaceTx | Anndata | Giotto | SpatialFeatureExperiment | GEM/GEF+ adata |
| Programming language | Python | Python | Python/R | Python/ nextflow | Python | Python | Python | R | R/Python | pytho |
| Image-level QC | Yes | No | For IMC | No | Yes | Yes | No | No | No | no |
| Cell-level QC | Yes | Yes | For Imc | Yes | No | limited | Yes | Yes | Yes | yes |
| Gene-level QC | Yes | No | For IMC | limited | No | Yes (at spot level) | No | No | Yes | no |
| Dimensionality reduction + clustering | Yes | Yes | Yes | No | Yes | No | Yes | Yes | Yes | yes |
| Tunability | Yes | Limited | No | No | Yes | No | Yes | Yes | Yes | No |
| Usability | CLI, API, GUI | CLI, API | API,CLI | Nextflow CLI | API | CLI (CWL?) | API | API | API | API |
| Normalization | tunable | Lib size | For IMC | No | intensities | No | Lib size | Yes, multiple | logNorm | Lib size |
| Integration | scVerse | scVerse | Bioconductor | no | None | None | scVerse | Giotto | Bioconductor | no |

**Supplementary table 1: overview of open-source Spatial Transcriptomics Pipelines**
